# Initial refinement of data from video-based single-cell tracking

**DOI:** 10.1101/2022.04.26.489486

**Authors:** Mónica Suárez Korsnes, Reinert Korsnes

**Affiliations:** Norwegian University of Science and Technology (NTNU), Department of Clinical and Molecular Medicine, NO-7491 Trondheim, Norway; Korsnes Biocomputing (KoBio), Trondheim, Norway

**Keywords:** single-cell tracking, phenotypic signature, big data, cancer diagnostic methods, daughter cells

## Abstract

**Background:** Video recording of cells offers a straightforward way to gain valuable information from their response to treatments. An indispensable step in obtaining such information involves tracking individual cells from the recorded data. A subsequent step is reducing such data to represent essential biological information. This can help to compare various single-cell tracking data providing a novel source of information. The vast array of potential data sources highlights the significance of methodologies prioritizing simplicity, robustness, transparency, affordability, sensor independence, and freedom from reliance on specific software or online services.

**Methods:** The provided data presents single-cell tracking of clonal (A549) cells as they grow in two-dimensional (2D) monolayers over 94 hours, spanning several cell cycles. The cells are exposed to three different concentrations of yessotoxin (YTX). The data treatments showcase the parametrization of population growth curves, as well as other statistical descriptions. These include the temporal development of cell speed in family trees with and without cell death, correlations between sister cells, single-cell average displacements, and the study of clustering tendencies.

**Results:** Various statistics obtained from single-cell tracking reveal patterns suitable for data compression and parametrization. These statistics encompass essential aspects such as cell division, movements, and mutual information between sister cells.

**Conclusion:** This work presents practical examples that highlight the abundant potential information within large sets of single-cell tracking data. Data reduction is crucial in the process of acquiring such information which can be relevant for phenotypic drug discovery and therapeutics, extending beyond standardized procedures. Conducting meaningful big data analysis typically necessitates a substantial amount of data, which can stem from standalone case studies as an initial foundation.

## 1 Introduction

The goal of this contribution is to showcase the potential benefits of sharing refined single-cell tracking data obtained from video recordings. Recent advances in single-cell research make it more interesting to follow individual cells over time to gain dynamic information from them at the individual level [Skylaki et al., 2016]. Data from such observations can reflect several processes and signalling pathways inside cells and between them. Tracking single cells in video aspires to provide this type of information, which can already be lost when working on fixed dead cells. Such tracking can also contribute to characterize phenotypic states and quantify them, as permanent or temporary [Tata and Rajagopal, 2016, Gupta et al., 2019]. It can provide data on lineage relationships between cells and their descendants, contributing to trace population dynamics and insight into possible pathological outcomes [Woodworth et al., 2017].

Single-cell tracking is especially relevant for studying cancer cells, which are known to exhibit highly adaptable behavior during treatments. Cancer cells can rapidly alter their gene expression profiles to adapt to new microenvironments, making them difficult to target effectively [Lüönd et al., 2021]. This high plasticity also enables cancer cells to fuse during close cellular interactions, generating hybrid subpopulations with enhanced tumorigenicity and metastatic capacity [Melzer et al., 2018, Shabo et al., 2020, Hass et al., 2020]. In addition, cancer cells can display significant phenotypic heterogeneity within genetically identical populations as a result of unique transcriptomes and proteomes [Chang et al., 2008]. This heterogeneity, which can be driven by epigenetic alterations, poses a challenge for guiding personalized treatments [Wakita et al., 2013, Bheda and Schneider, 2014, Shinjo and Kondo, 2015, Bintu et al., 2016]. Tracking single cells over time can provide valuable insights into lineage relationships and population dynamics, shedding light on the mechanisms behind these phenomena.

Several authors emphasize that single-cell tracking from video has broadened the spectrum in mammalian signalling networks, drug development and cancer research [Regot et al., 2014, Suman et al., 2016, Van Valen et al., 2016, Koh et al., 2017, DuChez, 2018, Korsnes and Korsnes, 2018, Emami et al., 2020, Fujimoto et al., 2020, Fazeli et al., 2020, Ghannoum et al., 2021]. Korsnes and Korsnes [2015, 2017, 2018] showed statistics from systematic single-cell tracking during several days, elucidating heterogeneous cell response and induction of cell death mechanisms. This tracking also allowed detection of inheritable traits, such as vacuolar transfer from mother to daughter cells. Inheritance may here be significant for the interpretation of observations related to autophagy signalling [Korsnes et al., 2016, Klionsky et al., 2021].

Andrei et al. [2020] pointed out different types of observables from tracking two-dimensional (2D) cell cultures and that might have biological relevance in cellular studies. 2D cultures have provided a wealth of information on fundamental biological process and diseases over the past decades [Langhans, 2018]. The advantage to use these models for tracking single cells is their low cost and reproducibility as compared to three-dimensional platforms [Nishida-Aoki and Gujral, 2019, Edlund et al., 2021, Helgadottir et al., 2021]. Two-dimensional models can easily integrate subsequent biochemical analysis and act as surrogate measurements for the 3D situations [Capuzzo and Vigo, 2021].

Three-dimensional (3D) models are under active development to better represent the complexity of living organisms during *in vitro* research [Yong et al., 2017, Finnberg et al., 2017, Puls et al., 2017, Fontana et al., 2021]. However, they still do not recapitulate micro-environmental factors, being only reductionist of the *in vivo* counterpart [Langhans, 2018, Capuzzo and Vigo, 2021]. 3D cell culture models are currently application specific and experiments with them are difficult to check for repeatability [Langhans, 2018]. Current 3D platforms do not allow acquisition of cellular kinetics with a high spatial and temporal resolution over a long period of time [Wen et al., 2021, Capuzzo and Vigo, 2021]. High-content screening (HCS) platforms are emerging, however visualization of 3D structures growing within complex geometrical structures remain still a big challenge mainly due to optical light scattering, light absorption and poor light penetration with prolonged imaging acquisition times [Langhans, 2018]. Microfluidic devices under highly controllable environmental conditions is a well-established operation in ongoing research Schmitz et al. [2021]. However, optimal nutrient supply and sufficient cell retention, especially for the long-term cultivation of slow growing cells as well as motile cells still requires a reliable cell retention concept to prevent permanent cell loss, which otherwise compromises qualitative and quantitative cell studies [Schmitz et al., 2023].

This study restricts to processing data from tracking individual cells growing in two-dimensional (2D) monolayers. The intention is to show, by simple examples, the potential utility of large collections of such data, allowing users to compare their experiments with many previous similar experiments. Such collections would facilitate big data analysis, taking advantage of weak correlations in large amounts of data. The source of these data may be video recordings of diverse quality, assumed as by-products from experiments worldwide. The present example data therefore, for the sake of simplicity, only represent positions (tracks) of individual cells and their eventual division and death during recording. They originate from previous work on Yessotoxin (YTX) [Korsnes and Korsnes, 2018]. This small molecule compound can induce different cell death modalities [Korsnes, 2012]. The broad spectrum of cellular response to YTX suits the present illustrations. The richness of responses from it may also make the compound an interesting candidate to probe cells for properties.

Data collections from single-cell tracking can be a resource for both experimental work and statistical investigations, including fault-tolerant big data analysis to search for patterns of biologic relevance. The present data processing may also have direct interest for processing videos aimed for special studies on possible emergence of rare or resistant subpopulations among cells subject to toxic agents, potential for metastasis or early screening for drug discovery. Another actual application is simply to check for the healthiness of cell populations, including testing for contamination.

The data analyses below relates cells in pedigree trees, where the initial cells are the ancestors (roots). These trees facilitate classification of cells in subpopulations according to a combined analysis of the cells in each tree. An example of such a combined analysis is to count the number of dying cells in each pedigree tree. The statistics below apply this simple idea assuming that cells in pedigree trees, with no cell death, might define a special resistant subpopulation. It reflects, for different subpopulations, variation in cell speed, correlations between sister cells as well as relocation and tendency of clustering. The authors conjecture that such data summaries can guide computerized search after patterns and causal relations in large sets of single-cell tracking data. The final proof of concept depends on access to such data sets.

A variety of relatively low-cost equipment apply to perform video-based single-cell tracking in 2D cellular models. Researchers can now in their most cost-effective way produce videos of living cells for subsequent analysis by remote (Internet/cloud based) services, as recently developed by Korsnes Biocomputing (KoBio)^1^. They may also do similar analysis/tracking using their own favorite tools, such as Image J/ TrackMate [Tinevez et al., 2017]. The supplementary data illustrates the potential transparency and software/equipment independence of such data production^2^ facilitating sample inspection. Perturbation of data values can reveal if analysis results are sensitive to measurement errors. These factors make such data relevant for contribution to biological databases reviewed by Zou et al. [2015], Haniffa et al. [2021] and Osumi-Sutherland et al. [2021]. The main intention here is taking advantage to utilize data from simple and low cost recordings to create synergistic value from sharing data on cellular behaviour.

## 2 Materials and Methods

### 2.1 Toxin

Yessotoxin (YTX) was obtained from the Cawthron institute (Nelson, New Zealand). YTX was dissolved in methanol as a 50 μm stock solution. The stock solution was after diluted in RPMI medium (Lonza, Norway), achieving a final concentration of 2 μm YTX in 0.2 % methanol. Treated cells were incubated with 200 nm, 500 nm and 1000 nm YTX and control cells were incubated with 0.2 % methanol as vehicle.

### 2.2 Cell culture

A549 cell lines were provided by Dr. Yvonne Andersson and Dr. Gunhild Mari Mœlandsmo from the Institute of Cancer Research at the Norwegian Radium Hospital. Cells were cultured in RPMI 1640 (Lonza, Norway), supplemented with 9 % heat inactivated fetal calf serum (FCS, Bionordika, Norway), 0.02 M Hepes buffer 1M in 0.85 % NaCl (Cambrex no 0750, #BE17-737G) and 10 ml 1X Glutamax (100X, Gibco #35050-038), 5 ml in 500 ml medium. Cells were maintained at 37 °C in a humidified 5 % CO_2_ atmosphere.

### 2.3 Single live-cell imaging and tracking

A549 cells were plated onto 96 multi-well black microplates (Greiner Bio-One GmbH, Germany) for timelapse imaging. Cells were imaged into Cytation 5 Cell Imaging Reader (Biotek, USA), with temperature and gas control set to 37 °C and 5 % CO_2_ atmosphere, respectively. Sequential imaging of each well was taken using 10*×* objective.

The bright and the phase contrast imaging channel was used for image recording. Two times, two partly overlapping images were stitched together to form images of appropriate size. A continuous kinetic procedure was chosen where imaging was carried out with each designated well within an interval of 6 min for an 94 h incubation period. Exposed cells were recorded simultaneously subject to three different concentrations of YTX 200 nm, 500 nm and 1000 nm.

The single-cell tracking in this work was performed using the in-house computer program *Kobio Celltrack*^3^. The present data derives from previous work on YTX [Korsnes and Korsnes, 2018].

## 3 Results

### 3.1 Single-cell tracking

Figure 1 illustrates production of input data for the present analysis. The coloured dots in Figure 1a represent individual cell positions during 94 h of recording. The tracking also provides data on cell division and dead. The actual tools for tracking are outside the scope of this study, which in principle could rely on data from any functional tracking system.

**Figure 1:**
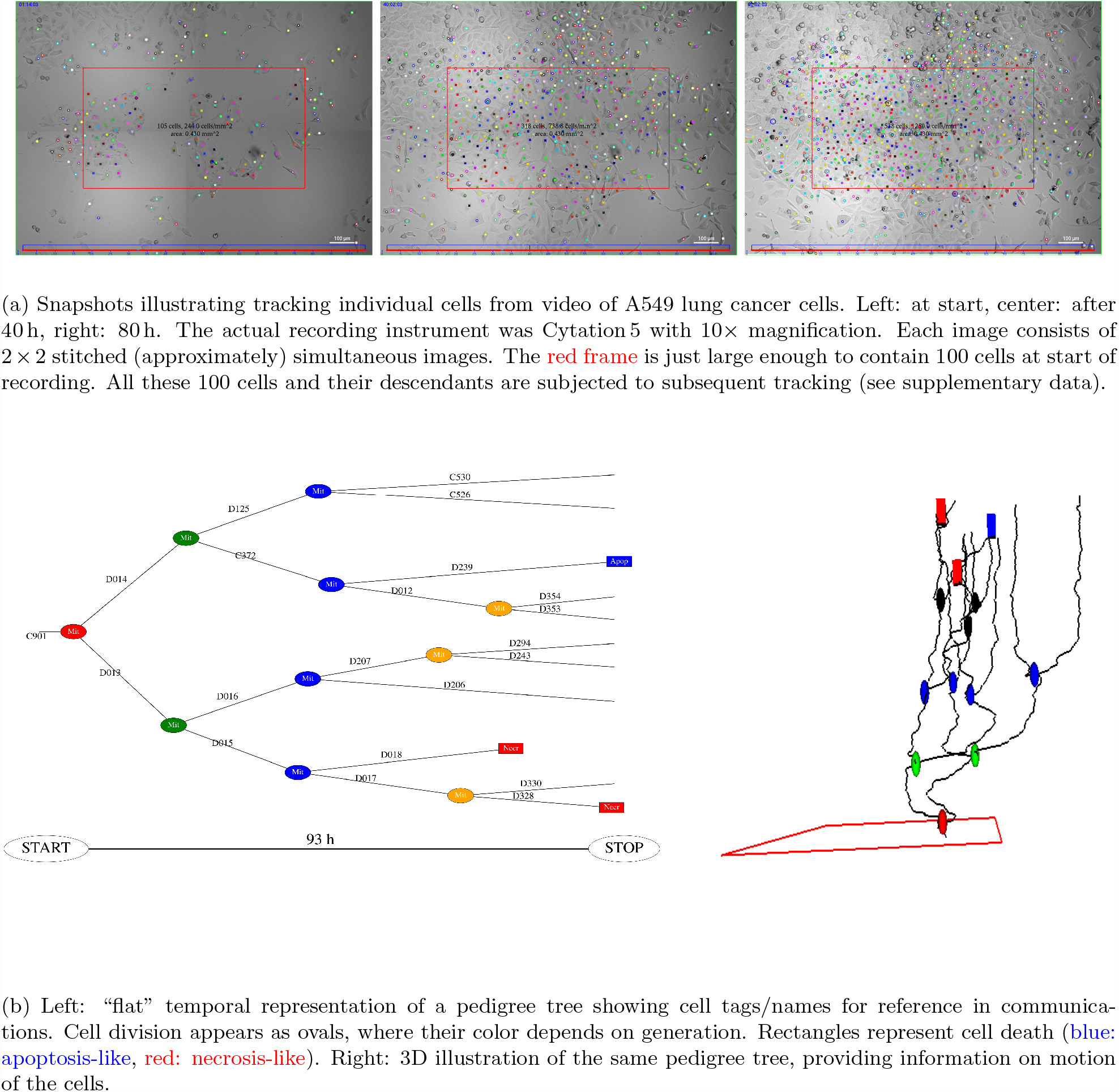
Illustration of production of single-cell tracking data for subsequent analysis and data compression aimed for big data analysis. Cells were in this case exposed to 200 nm YTX. The supplementary data includes video demonstrations of the actual tracking^2^.

Figure 1b illustrates data products from the prior single-cell tracking. The left part of the figure gives a time attributed graph representation of kinships between the descendants of a cell which is inside the red frame at start of recording. The right part illustrates the positions of these cells during recording. The horizontal positions (x-y coordinates) here represent spatial location and the height (z-coordinate) represents time. The red frame is here just large enough to contain 100 root cells at the start of recording. The present examples of statistical analysis is for the cells belonging to the pedigree trees starting inside such a red frame.

Figure 2 shows spatially located pedigree trees for cells exposed to YTX at three different concentrations. Cells in surviving lineages exposed to the highest YTX concentration (1000 nm) may appear to behave similar to cells subject to the lowest YTX (200 nm) concentration. It could reflect a resistant subpopulation.

**Figure 2:**
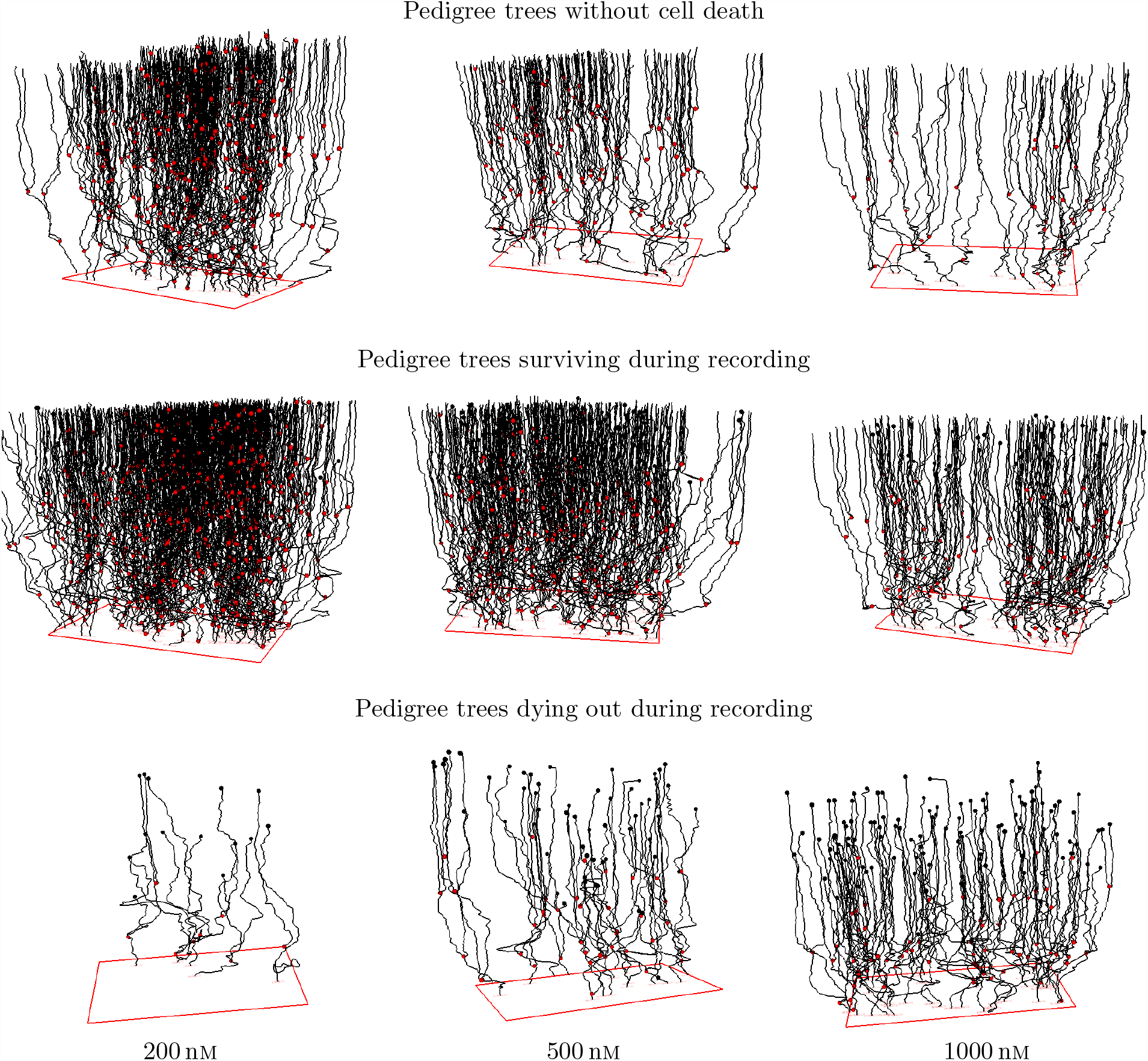
“Forest” of pedigree trees from tracking A549 cells exposed to YTX at concentrations 200 nm, 500 nm and 1000 nm. The upper row shows trajectories for cells in lineages without death (“resistant cells”). The middle row is for trajectories of cells in lineages where at least one cell live at the end of recording (“surviving pedigree trees”). The lower row shows trajectories for cells in lineages dying out during recording. Red and black dots represent cell division and cell death respectively. Note that single-cell tracking can provide more precise information on cell viability as compared to traditional bulk assays. These type of measurements are prone to overestimate cell survival due to prior apoptotic cell clearance and disintegration.

### 3.2 Single-cell viability

Cell tracking offers valuable insights into fundamental cellular properties like survival and proliferation, making it a crucial tool across various domains of cell research such as risk assessment for toxic agents, drug screening, and cancer research. Researchers studying the impact of specific toxic agents on a particular group of cells can enhance their understanding by comparing their findings with data from similar experiments conducted elsewhere. Efficient reduction of such data plays a pivotal role in facilitating meaningful comparisons and enabling access to relevant information within extensive data collections. This section presents prototypes of data reduction techniques aimed at achieving these objectives.

Figure 3a shows the change in size of distinct cell subpopulations during video recording. The graphs show the development of number of cells in pedigree trees with roots (initial ancestors) inside a frame centered in the video and just large enough to contain 100 cells at the start of recording. The population of cells belonging to the largest pedigree trees naturally grows faster than the total population. These cells potentially dominate in number after some time, if they inherit their tendency of cell division and survival. Correlations between proliferation and survival of descendants of sister cells (see Figure 4) can indicate such inheritance.

**Figure 3:**
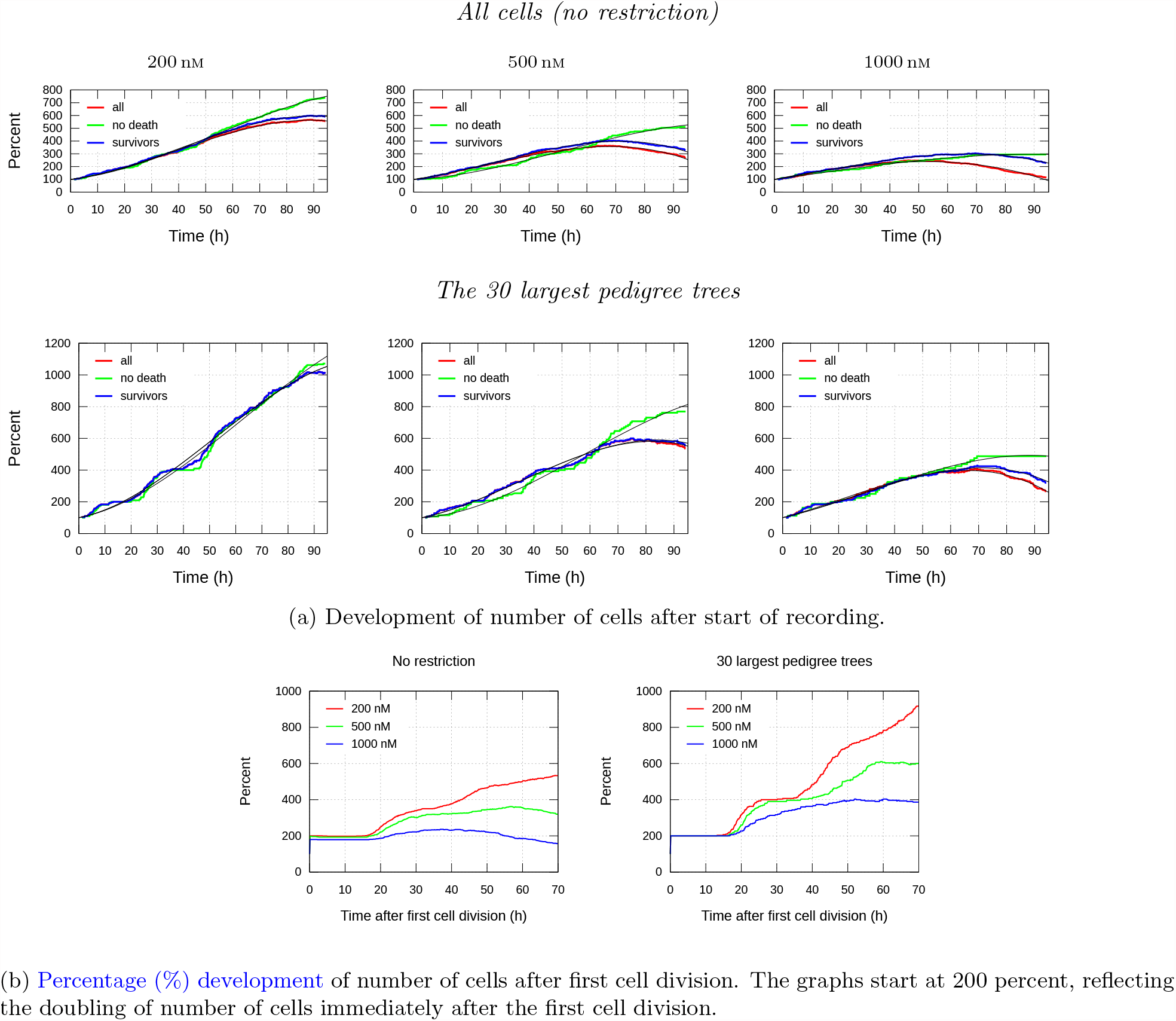
Illustration of different views of cell proliferation for A549 cells exposed to 200, 500 and 1000 nm YTX concentrations. Note that “*all*” refers to all cells in the red frame (see Figure 1a); “*no death*”: cells in lineages with no death; “*survivors*”: cells in pedigree trees where at least one cell lives at the end of recording. Note the smoothness of the graphs, enabling effective parametrization (“data compression”).

**Figure 4:**
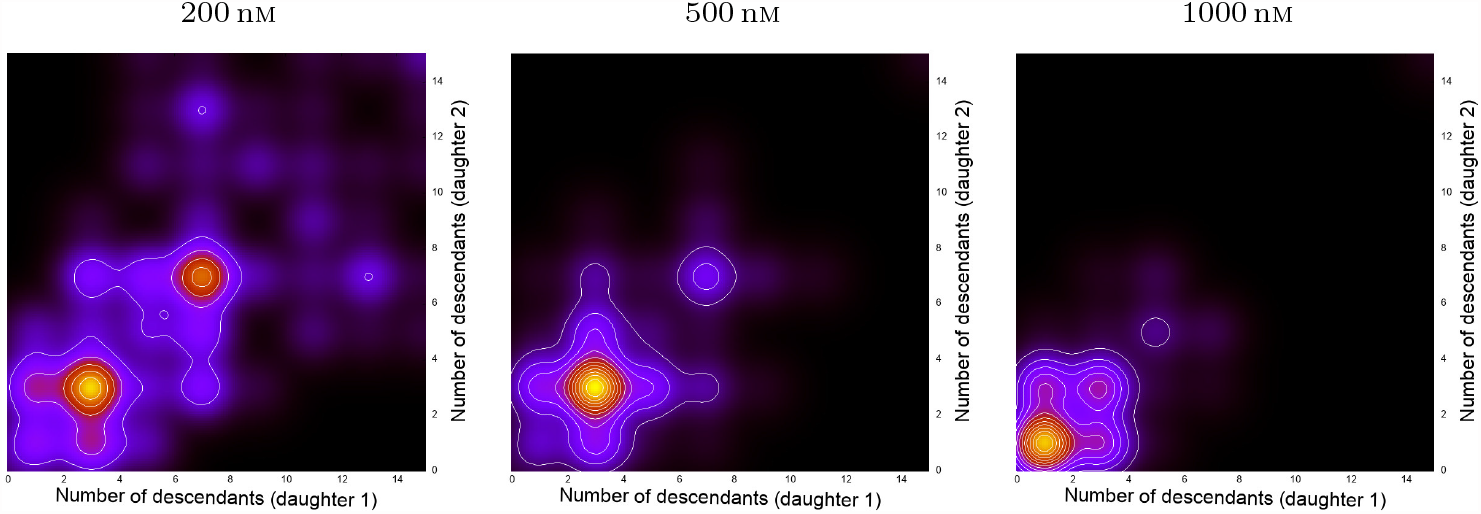
Results from kernel smoothing (bandwidth 1.5) of stack plots for number of descendants of first generation sister cells within 70 h after their birth. The plots are for cells born within 20 h after start of recording. The two apparent clusters in the plot for cells exposed to 200 nm YTX indicate inheritance from the common mother cell.

The lower row in Figure 3a illustrates that cell “viability analysis” based on single-cell tracking can provide information beyond results from traditional bulk analysis. The black solid lines in Figure 3a represent a third degree polynomial model fit to the data:

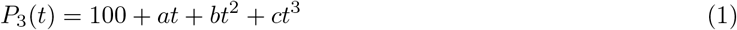

where *a, b* and *c* are the (model) parameters and *t* represents time. Polynomials (or Taylor expansions) are generally a convenient way to represent smooth (“simple”) functions and to compress data (representing it by three parameters). Parameters from fitting a complex biologically justified model may not necessarily represent more biological relevant information if they are less effective to compress data.

Assume fitting a Taylor model (Equation 1) to data as above (see Figure 3a). Consider the resulting parameters as a point, ***P*** = (*a, b, c*), in the three-dimensional parameters space. Similar parameters from various experiments will give a set of points in the parameter space. If these points spread out close to for example a 2D structure (embedded in the 3D space), then there should, intuitively, be hope for finding statistical models with two parameters (instead of three) providing a biological interpretation/understanding. Voids in the parameter space can also represent knowledge.

Figure 3b shows percentage development of number of cells in pedigree trees as a function of time after the first cell division. The left part of the figure is for the all 100 pedigree tress (initiating in the red frame as explained above), and the right part is for the 30 largest pedigree trees. The figure shows that cells exposed to the lowest concentration of YTX (200 nm) tend to follow a regular timing for cell division, as opposed to those subject to the highest concentration (1000 nm). This tendency is most expressed for the largest pedigree trees (right part of the figure).

### 3.3 Speed

Measurements of cell speed offer valuable insights into cellular conditions following various treatments. This information holds significant prognostic value by providing indications of cellular response and potential outcomes associated with specific interventions. For instance, it can contribute to the identification of distinct migration and persistence values that may correlate with the rate of intravasation [Ghannoum et al., 2023].

Similar arguments for data reductions of viability, discussed in Chapter 3.2, are also applicable to cell speed. It is worth noting that viability and speed are likely to be correlated, which presents additional opportunities for data reduction, including dimensionality reduction techniques [Fodor, 2002].

Track length for a cell during a period of time *t* (divided by *t*) can intuitively define its average *speed* during that period. However, track length is not in practice directly available nor be it well-defined for imprecise and irregular positional data, where measures of length can depend on resolution. *Cell speed* could (ad hoc) refer to movements of a given defined point in a cell (for example, the mean point of the nucleus/nuclei). However, it may principally be looked at as a spatio-temporally localized (statistical) property of a cell. Future work may assume an “uncertainty principle” where a positional data point is considered as a random selection from a set of possible positions depending on the tracking method. An alternative approach is to increase the level of sophistication and replace the concept of “cell speed” with temporal change in the (segmented) set of points covered by an actual cell.

Estimates of positions are, for any definition, imprecise for low quality imagery data. This work therefore, for the sake of simplicity, demonstrates Gaussian kernel smoothing and interpolation [Wand and Jones, 1994] to define speed. The actual bandwidth is 15 min. Perturbations of estimates of cell positions may help to reveal how final results are sensitive to this choice of bandwidth. The authors leave out this exercise for a separate study. Note that big data approaches may in principle automatically sort out useful definitions of speed.

Figure 5 shows distributions of the eight hours centered moving generalized mean speed for cells in lineages with and without death during recording. The upper and third row are for the regular mean, whereas the second and lower row similarly show the fourth power mean for the same data. This example illustrates a possible data product that presumably could provide information to big data analysis. The power mean *M*_*p*_ is increasingly more sensitive to the highest speeds for increased values of *p*. The distribution for *M*_4_, for example, seems to be more sensitive to cell death in lineages as compared to lineages with no cell death. One can expect that the power mean *M*_*p*_ for *p* = 1, 2, …, *n* will in a compact way reflect the distribution of speed for a restricted value of *n*.

**Figure 5:**
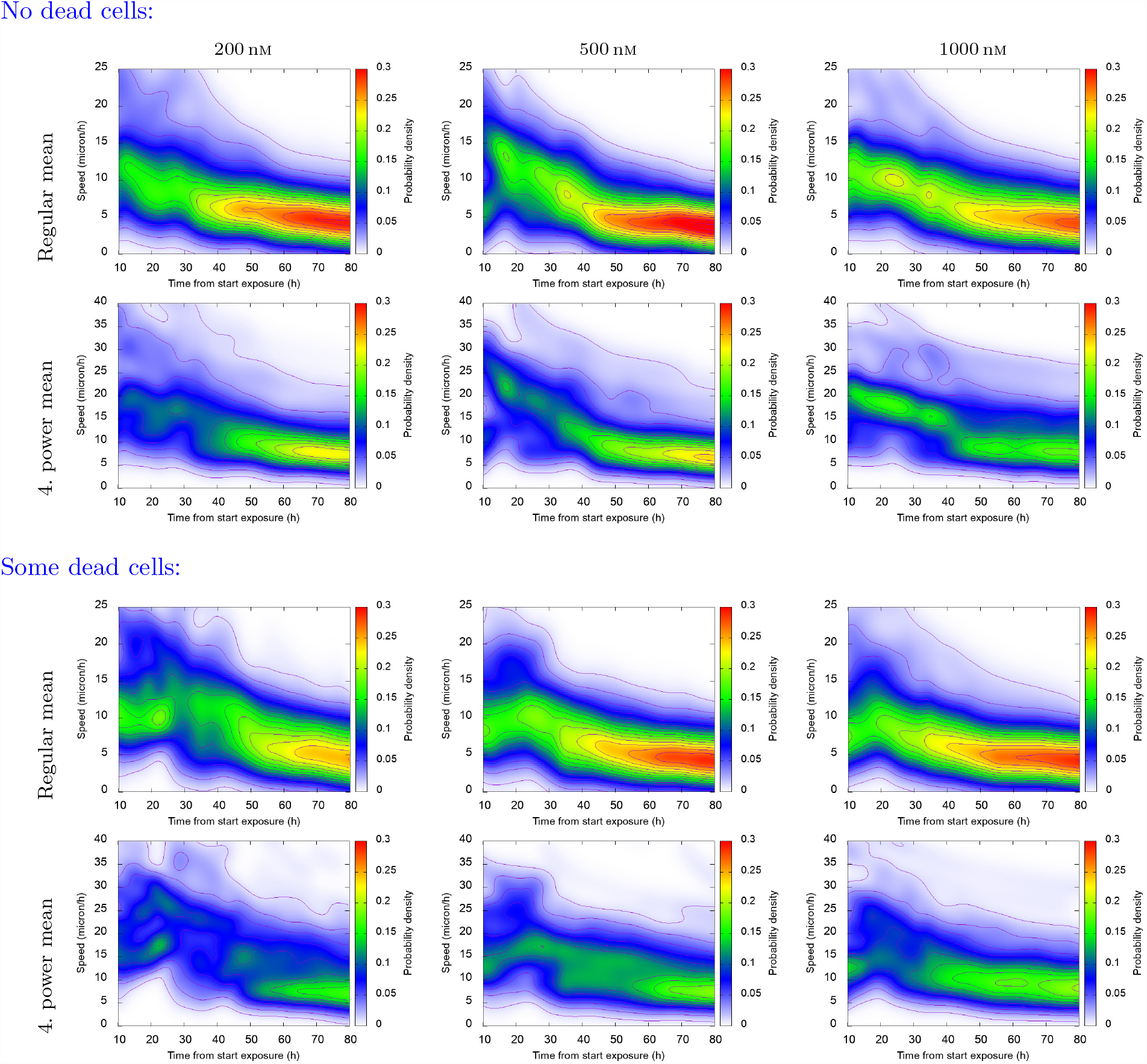
Distribution of 8 hours centered running generalized mean of speed of cells during recording. The top and third row show regular (first order) mean, and the second and fourth row show fourth power mean (*M*_*p*_, *p* = 4). Note the difference between the distributions, especially at the first part of the recording.

### 3.4 Correlation between descendants of sister cells

Correlating or analyzing the mutual information^4^ between parameters of sister cells can reveal signaling downstream lineages. The treatment of cells can affect their signaling and potentially introduce noise during cell division affecting the behaviour of descendant cells. As a result, single cell tracking data has the potential to capture and reflect this valuable information. When multiple cell types exhibit similar responses to similar treatments performed at different laboratories, it can provide deeper insights into cellular reactions. By comparing single cell tracking data from different experiments, we can facilitate the discovery of robust findings. This section outline ideas for summarizing or reducing the data to facilitate this search.

Figure 6 shows joint distributions for the total track length of first generation sister cells and their descendants 60 h after the birth of these (initial) sister cells. These statistics restrict to sister cells born within 30 h after start of recording. The estimates result from using the algorithm scipy.stats.gaussian kde from SciPy^5^ with defaults settings (i.e., the ‘scott’ method defines the estimator bandwidth). Section 3.3 outlines the present estimation of length from imprecise positional data (applying Gaussian kernel smoothing).

The joint distributions of Figure 6 show positive correlations and hence reflect inheritance from mother cells to their daughters. The authors will not further speculate on the biological significance of these statistics, since they only reflect results from one experiment. However, a main finding here is that such distributions are sensitive to cell treatment. One may therefore suspect such data summaries to be relevant for big data analysis. The regularity of such distributions enables effective parametrization (or data compression) to help search in large databases.

**Figure 6:**
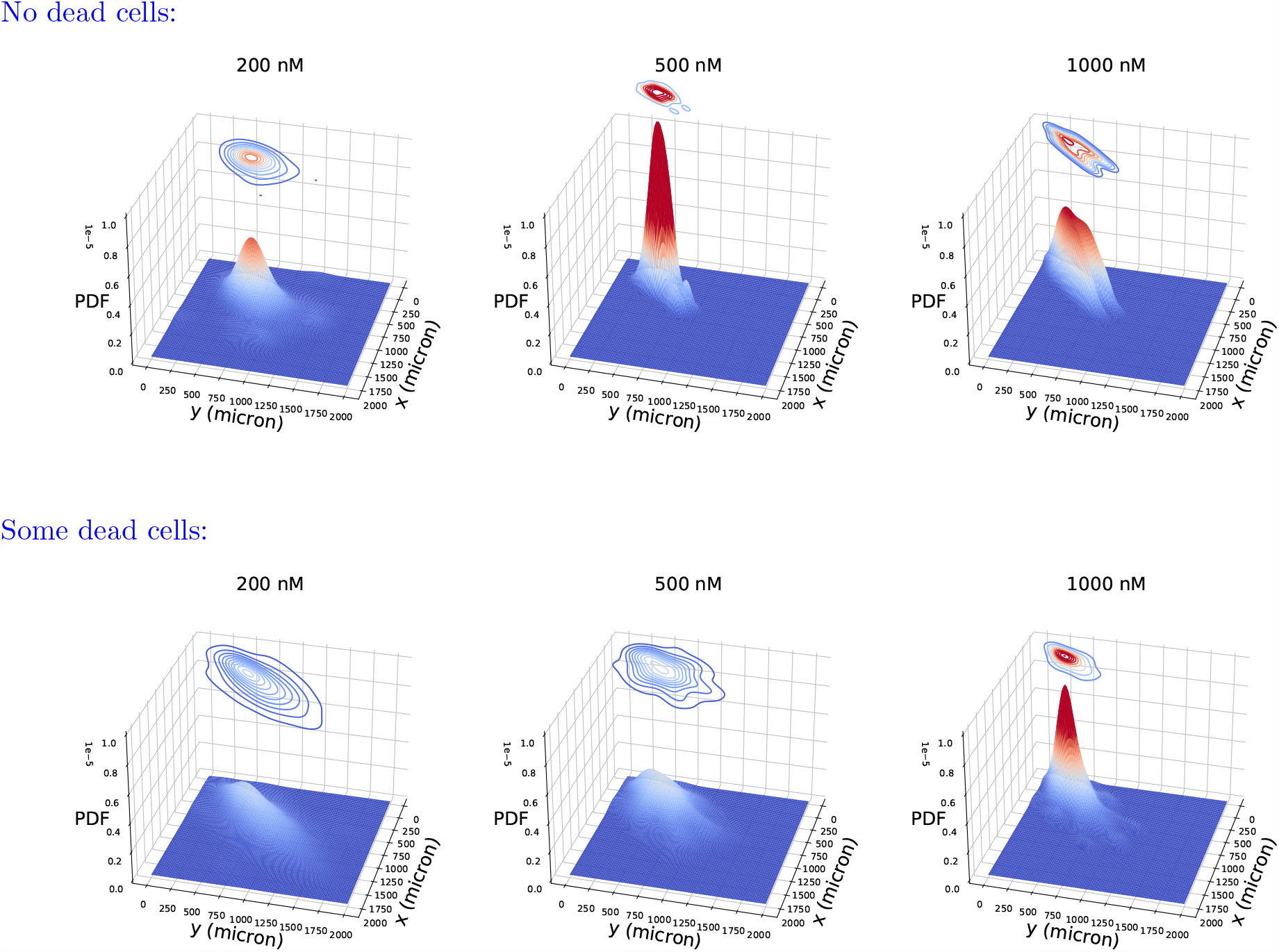
Joint probability density function (PDF) of total track length (*x* and *y*) for the first generation sister cells and their descendants 60 h after the birth of these (initial) sister cells. The cells are subject to YTX exposure at concentrations 200 nm, 500 nm and 1000 nm. The upper row shows distributions for the pedigree trees with no cell death, and the lower one shows pedigree trees with some cell death.

Figure 7 supports the notion of signaling downstream lineages by demonstrating visual evidence of morphological similarities among cells within the same pedigree tree, in contrast to the surrounding cells. Moreover, the corresponding pedigree trees and movements also exhibit resemblances. These observations strongly imply that establishing connections between cells in pedigree trees can significantly aid the analysis of single cells. Classification of cells, for example, often involves a certain level of uncertainty. However, by adopting a combined classification approach specifically designed for pedigree trees, it can be feasible to mitigate this inherent uncertainty.

**Figure 7:**
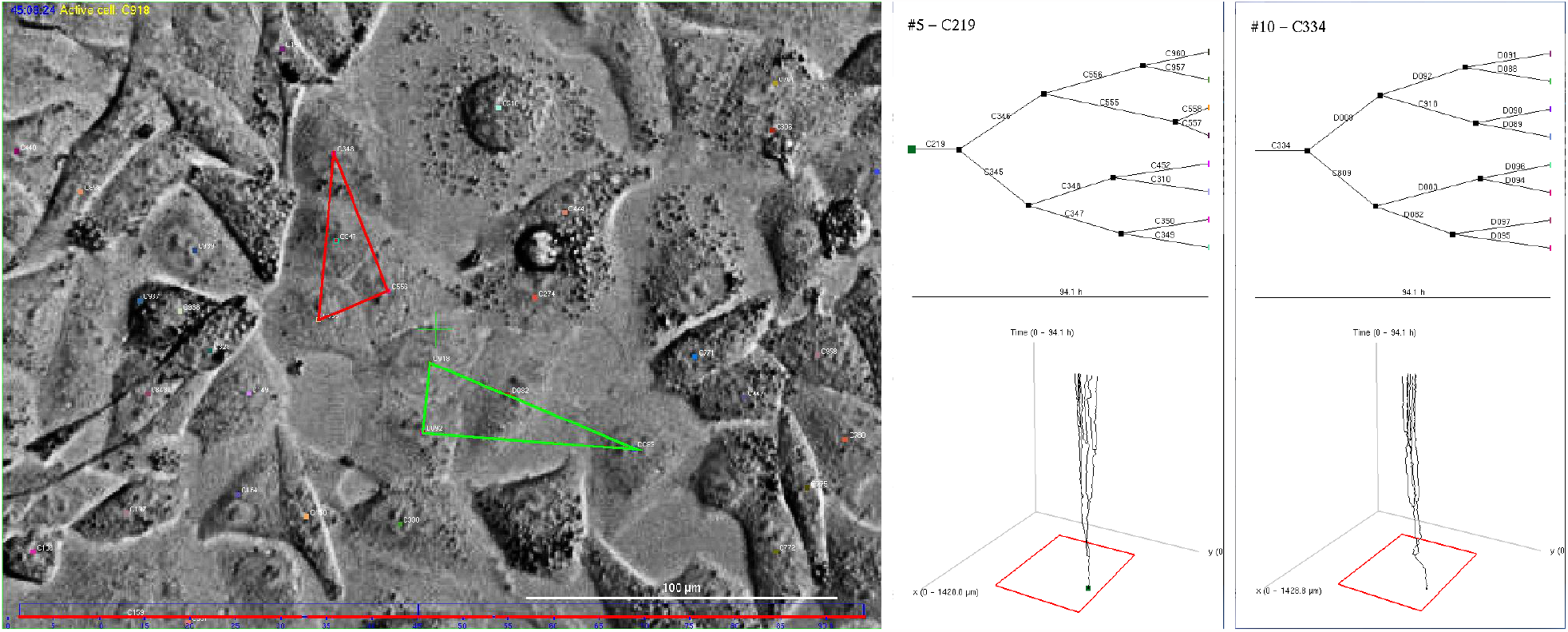
Visual illustration of morphological similarities between cells in the same lineage. This is an argument that combined analysis of cells in pedigree trees can provide more information as compared to analysis of cells without knowing their close relatives. Left: Snapshot from video of A549 cells after 45 h expossure to yessotoxin. The middle and right sections depict pedigree trees, with the lower portion demonstrating the movement of cells in the above pedigree tree during recording. The time axis is represented upwards. The red triangle points out cells in the pedigree tree with root cell C219 (middle of figure) whereas the green triangle points out cells in the pedigree tree with root cell C334 (right part of figure). Note that these cells form clusters.

### 3.5 Mean square displacement (MSD) of first generation daughter cells

The mean square displacement (MSD) of cells over time is a measure that captures both their speed and directional persistence. Statistics from it can presumably help big data analyses to find causal relations in large sets of single cell tracking data. Such data can also have direct interest in special studies. Ghannoum et al. [2023] for example, argue for the importance of acquiring such data for better understanding tumour growth rate and size.

This section explores potential methods for extracting features from such data, aligning with the principles discussed in sections 3.2, 3.3, and 3.4. Figure 8a illustrates the MSD of first generation daughter cells, depicting their displacement as a function of time since birth. The figure is for cells in pedigree trees, with and without cell death during recording.The mean square displacement (MSD) of cells over time reflects both their speed and movement patterns. This section explores potential methods for extracting features from such data, aligning with the principles discussed in sections 3.2, 3.3, and 3.4. Figure 8a illustrates the MSD of first generation daughter cells, depicting their displacement as a function of time since birth. The upper row here shows the tendency of cells to need extra time to start drifting from their place of birth. Processing of more data may reveal if this extra time can be considered a “phenotype” useful for search in data from many diverse experiments.

**Figure 8:**
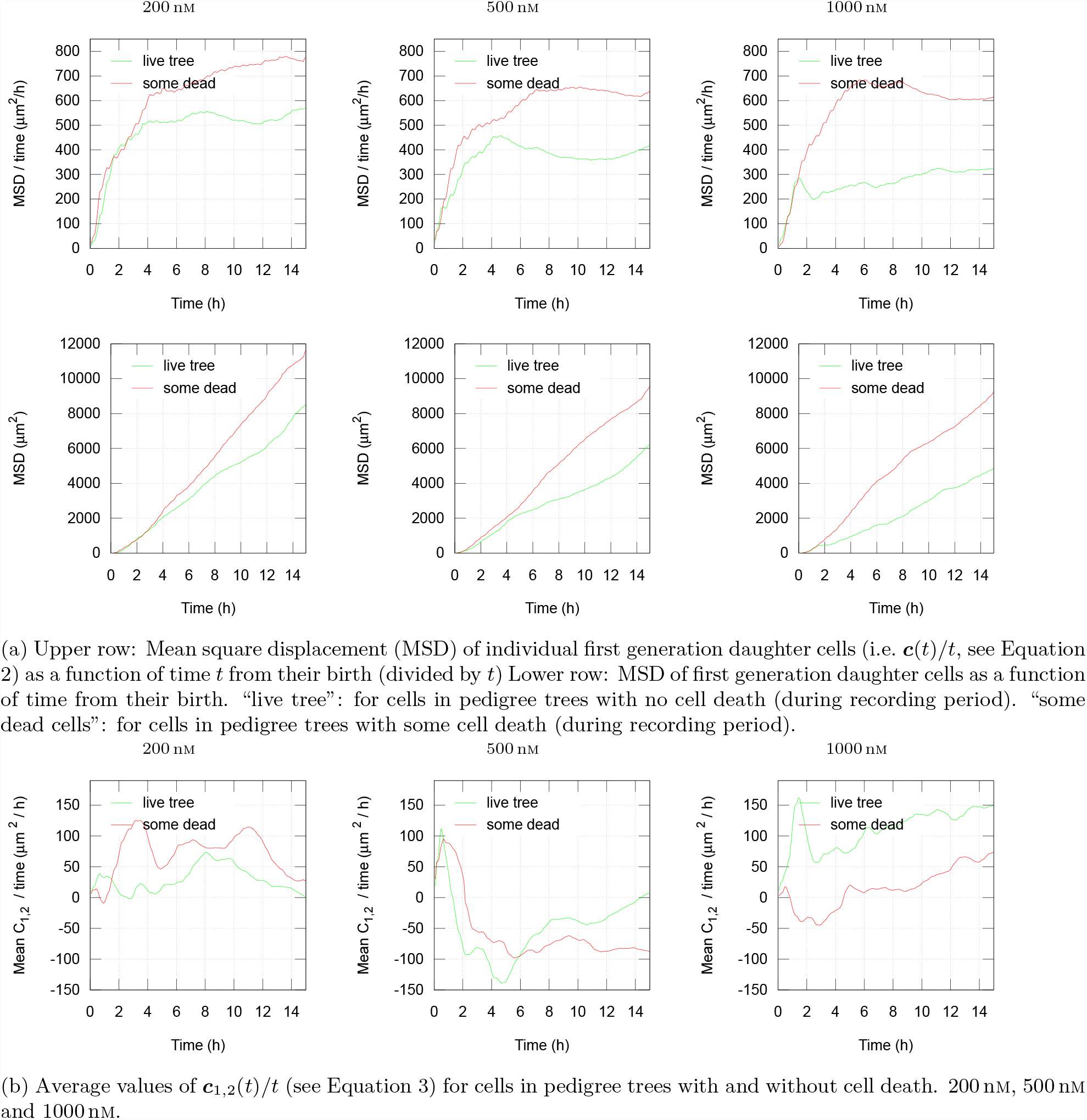
Examples of statistics of displacements of sister cells after their birth. The cells are subject to exposure by YTX at concentrations 200 nm, 500 nm and 1000 nm.

Note that Figure 8a indicates that cells in lineages with dying cells tend to move faster from their initial position as compared to cells with no observed cell death. A possible hypothesis is that cells with the strongest (inheritable) tendencies to move, are more vulnerable to the actual toxin (YTX) as compared to the others. One may also relate the observation to the concept of “fight-or-flight” reaction, where many types of cells respond to a variety of stressors in a reasonably standardized fashion, which allows them to combat the offending stimulus or escape from it [Goligorsky, 2001].

If the movement follows a “memory-less” Brownian type motion, the graphs for the upper row in Figure 8a would appear as straight horizontal lines, while the lower row would exhibit straight upward tilting lines. However, the actual graphs of Figure 8a reflect that the direction of movement tend to be independent of the direction about 4 h to 6 h earlier. The period up to about 4 h is “memory time” reflecting how long cells tend to keep their direction. It can partly correlate with cell shape, assuming elongated cells move in their longitudinal direction.

Assume the vector ***r***(*t*) represents the relocation of a cell *t* time units after its birth. The vector dot (inner) product

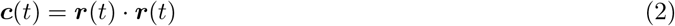

then gives this distance squared (equal |***r***(*t*)| ^2^). Figure 8a shows average values for *c*(*t*) for two subsets of cells where *t* ranges from 0 to 15 h. A tempting idea is slightly to modify this elaboration and check for average value of

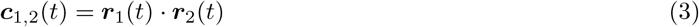

where ***r***_1_(*t*) and ***r***_2_(*t*) each represent the positions of a couple of siblings (sister cells) *t* time units after their birth. Figure 8b shows an example of results from such a numerical experiment. A motivation for this test is the conceptual simplicity and the pure formal similarity between the Equations 2 and 3. The authors have no specific biological interpretations of these graphs, except that they reflect the tendency for sister cells to follow each other after their birth. This tendency seems to depend on exposure.

### 3.6 Material exchange and trait inheritance

Moving cells are capable of maintaining close proximity for extended periods, which may suggest intercellular communication or material exchange that can impact their behavior. The specific characterization of this behavior is a subject for future research. Figure 9 exemplifies the identification of these events where cells exhibit prolonged closeness. This type of data may have special interest for co-culture or studies on differentiation where interactions are crucial. Cells can interact through physical contact, surface receptorligand interaction, cellular junctions, and secreted stimulus [Nishida-Aoki and Gujral, 2019]. Understanding these type of interactions can contribute to decipher the complex network of interaction between cells, helping to improve therapeutics [Nishida-Aoki and Gujral, 2019].

**Figure 9:**
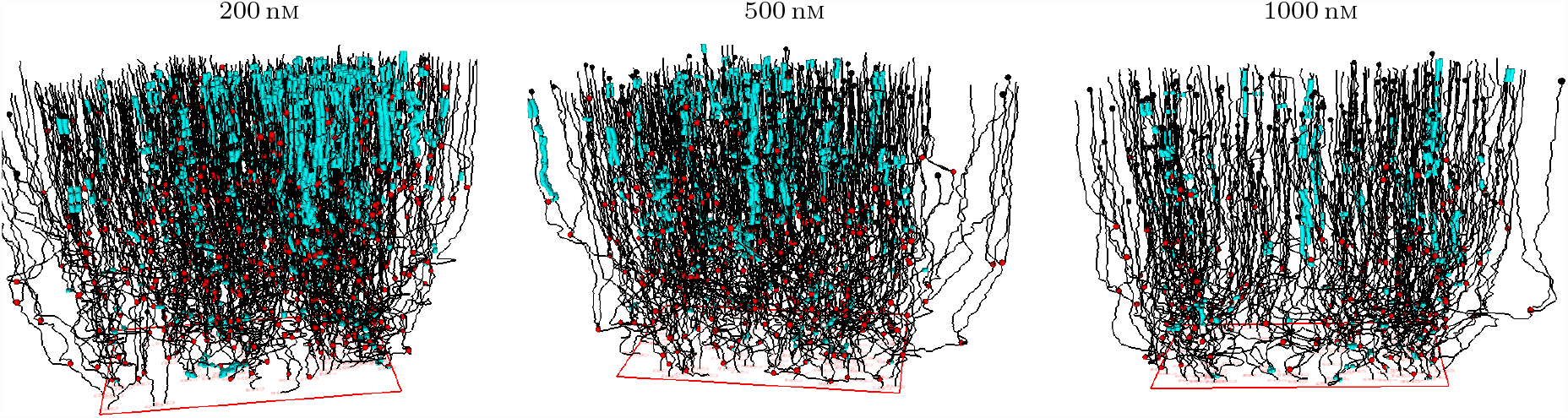
Forest of pedigree trees including identification of events where cells stay at the vicinity of each other for at least 4 hours 2 hours apart from their birth (cell division). The cells were subject to YTX exposure at concentrations 200 nm, 500 nm and 1000 nm.

Analyses of “forests” of pedigree trees can reflect effects from events where cells absorb debris from dead cells and transfer it to their descendants. Figure 10 shows an example of such behavior where a cell includes an apoptotic body from a neighboring dying cell. Such apoptotic bodies can subsequently appear as vacuoles in the absorbing cell. Sets of such vacuoles in a cell are traceable throughout cell division by comparing their size and number.

**Figure 10:**
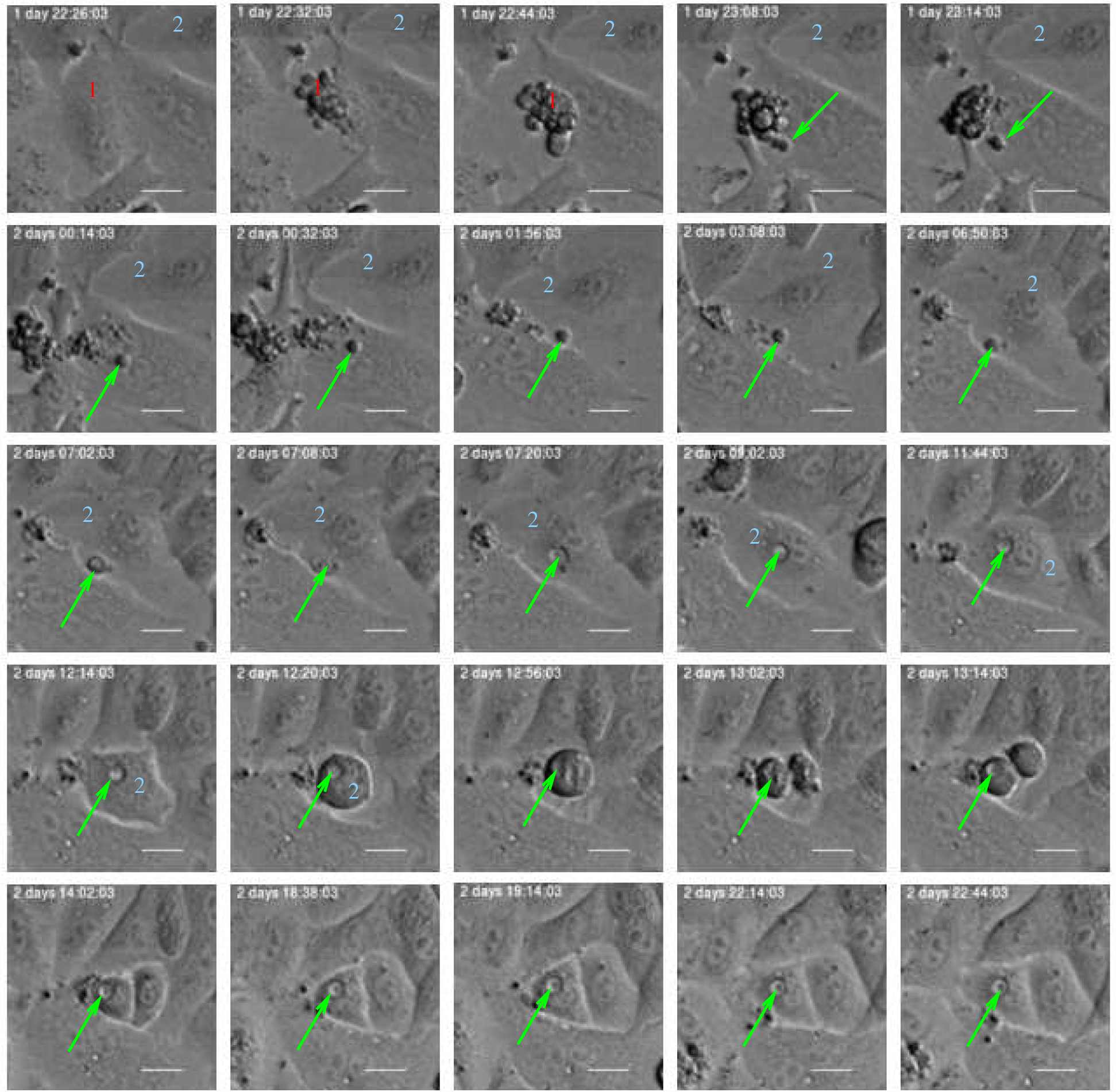
Example where an apoptotic body (green arrow) from a dying cell (1) ends up as a vacuole in a neighboring cell (2) which subsequently divides, and the vacuole ends up in one of the daughter cells. Detailed inspection of many cells in video can reveal such rare events and shed light on epigenetic heritage and generally signalling downstream pedigree trees. Such signalling is an argument to study lineages as independent entities and for example apply information on lineage relations when for example classifying cells.

## 4 Discussion

This work illustrates a number of possible methods to refine (or compress) data from video-based singlecell tracking. The main intention is to provide relevant input for big data analysis (or machine learning in general) to identify biomarkers for better diagnosis and prognosis. Well proven fault-tolerant computerized methods are here available to search for causal relations in large data sets [Theodoridis, 2015, Kotsiantis et al., 2007, Linero, 2017]. The principle of Occam’s razor [Domingos, 1998, 1999] can guide the search, favouring simplifications and approximations. It can be considered as a contradiction to anticipate the exact result from trying big data analysis methods, nor can one expect to anticipate what refinement methods are most effective. Successful big data analysis is (similar to data mining) assumed beyond the reach of human brains. However, their result may finally be understood by humans.

Big data methods go beyond assuming linear association between variables. The present examples therefore restrict to visual/intuitive illustrations of data refinement left for further processing. Existence of several local maxima in joint distributions (clustering) may, as an example, reflect significant biological information. The left part of Figure 4 illustrates this point. It shows two main maxima of the joint distribution of number of descendants of sister cells. This may indicate inheritance of robustness/viability making it likely for the most robust cells finally to dominate in number (which could be relevant for prognoses in cancer).

The present examples of refinement methods typically show different behaviour of cells in pedigree trees with cell death as compared to the behaviour of cells in pedigree trees without cell death (during recording). Some of these examples also show correlations between sister cells or descendants of sister cells. This is an argument to treat whole pedigree trees as individual entities in the initial data refinement.

Successful application of big data analysis can, in addition to sort out causal relations, give the possibility to search for similarities between the behavior of cells in many experiments. Methods to compare experiments can in general be an important part of a collective knowledge base of cell behavior.

Recent progress in techniques for sparse representations, compressive sensing and machine learning [see e.g. Papyan et al., 2018, Mousavi et al., 2019] give a perspective of direct automatic identification of actual biomarkers directly from video of cells. The present work contributes to this development by demonstrating initial refinement of data from single-cell tracking. These data summaries may also be of direct biological or medical interest in the conceptual frame of standalone experiments. They may in addition help the development of formal mathematical methods applying concepts from statistical physics [Banigan, 2013]. However, note that machine search for causality in data may utilize weak correlations without any immediate intuitive meaning.

This work illustrates derivation of the following parameters from single-cell tracking data which represent positions of individual live cells, their division, and death during several cell cycles:

- Number of cells in different classes of pedigree trees during video recording (Section 3.2). It may reflect that some pedigree trees consists of specially viable and resilient cells. This property seems to be already written into the root (ancestor) cell. Intrusive single cell analysis after tracking, while preserving track identities, may clarify corresponding mechanisms behind this resilience.
- Parameters from (representations of) speed distributions for various subsets of cells during tracking (Figure 5). The regularity of these distributions allow representations by few parameters (so-called sparse representation).
- Parameters from joint distributions of the size of (pedigree) subtrees for the first generation sister cells where they are roots cells (Figure 6). Such distributions can be parameterized by correlation coefficients, covariance, and shape parameters (or sparse representations).
- “Memory” time of trajectories for cells in subpopulations. Figure 8a reveals that cell trajectories can have a tendency to keep their direction, typically during 2 h to 4 h. This tendency can reflect cell shape.
- Tendency for cells to stay close to each other for periods. Figure 9 visualizes an example where cells tend to stay close for periods of time. Such events can potentially reflect intercellular communication and material exchange (see Figure 10). The tendency may have special interest for studies where communication between different cell types play a role. Tracking of cells in co-culture can in this case help to reveal how to affect such behavior.

An intention behind the present work is, as pointed out above, to promote ideas for better and easier comparison between different experiments. This would promote securing reproducibility of observations, which has emerged as a main concern in life science research in recent years [Hirsch and Schildknecht, 2019]. Easy exchange of raw and refined data is paramount in such quality assurance. Experiments on cells can include video recording of them under standard (common) conditions, and statistics from tracking the cells can reveal differences between experiments and which can affect their reproducibility. Tracking under standard conditions may in general reveal effects on cells and which otherwise may pass under the radar using bulk assays. This is an example of direct use of the present type of statistics.

Large-scale sharing of data from tracking single cells in video will naturally raise questions on robustness of results from initial analysis of them. Cells in different experiments may never be treated exactly the same way. Cells can be sensitive to photo-toxicity as well as possible molecular probes. Type of extracellular matrices and their proteins can also affect cellular behavior in test wells Vigilante et al. [2019]. Data analysis can reveal to what degree comparison of data from them still apply. It will be important to identify ranges of conditions for cells in which they will behave in comparable ways. It will also be important to identify conditions/treatments where cellular behavior is sensitive to small and uncontrollable perturbations. Data analysis may also reveal possible probabilistic views of results from observing cellular behavior.

Further development of sensors and software will extend the above restriction to data on cell positions, division, and dead. This will advance exploitation of its potential utility, as indicated by several authors [Van Valen et al., 2016, Tsai et al., 2019, Moen et al., 2019, Liu et al., 2020, Ghannoum et al., 2021]. Single-cell tracking from high quality imagery allows collecting data on phenotypical changes, otherwise difficult to measure from an end-point measurement such as single-cell RNA-sequencing (scRNA-seq) [Liu et al., 2020]. Furthermore, epigenetic states, protein expression and enzyme activity, can not only be inferred from changes in gene expression [Liu et al., 2020, Zhang et al., 2020]. Integrating single-cell tracking with RNA-seq analysis can therefore complement characterization of biological process by combining analysis of cellular phenotypes with gene expression profiles [Lane et al., 2017, Yuan et al., 2018]. These analyses allow overlaying phenotypic cell identity with genetic lineage information for a more comprehensive view of clonal relationships, since gene expression alone is not sufficient to classify cell states [Woodworth et al., 2017, Gerbin et al., 2021]. Integrating such analysis into cell ontology can help to discover a large variety of novel cell populations [Osumi-Sutherland et al., 2021]. Tracking individual cells can therefore complement current cell ontology efforts.

Big data analysis relies on a significant amount of data to derive meaningful insights, and accurately assessing the value of a dataset is only possible once it becomes available for analysis. As a result, the authors assert that a comprehensive roadmap for substantial and meaningful data sharing should involve the prototyping of statistical parameters and the creation of value through the execution of complementary specialized studies. The authors’ current contribution aims to serve as an inspiration for such specialized studies, motivating researchers to delve further into this field of research.

## Authors contribution

**Mónica Suárez Korsnes**: Conceptualization (equal); data curation (supporting); formal analysis (supporting); investigation (equal); methodology (equal); project administration (lead); software (supporting); validation (supporting); visualization (supporting); writing—original draft (equal); writing—review and editing (equal). **Reinert Korsnes**: Conceptualization (equal); data curation (lead); formal analysis (lead); investigation (equal); methodology (equal); project administration (supporting); software (lead); validation (lead); visualization (lead); writing—original draft (equal); writing—review and editing (equal).

## Acknowledgements

This work was supported by The Norwegian University of Science and technology (NTNU), Department of Clinical and Molecular Medicine.

## Footnotes

1 https://www.korsnesbiocomputing.no/

2 Supplementary data are available via https://user.korsnesbiocomputing.no (user *iniref_2022*, password korsnes1)

3 https://www.korsnesbiocomputing.no/

4 http://www.scholarpedia.org/article/Mutual_information

5 https://scipy.org/

